# Improving the monitoring of the invasive Blue Crab (*Callinectes sapidus*): combining environmental DNA and citizen observations

**DOI:** 10.1101/2024.10.24.620022

**Authors:** Erwan Delrieu-Trottin, Mathieu Bianic, Claude Miaud, Manon Garcia, Véronique Arnal, Régis Hocdé, Christophe Cochet, Nathalie Barre, Laetitia Cornil, Stéphanie Manel

## Abstract

1. Early detection is a crucial tool for identifying the spread of invasive species.
2. In this study, we validated a probe-based quantitative polymerase chain reaction (qPCR) assay for the detection of the invasive Blue Crab, *Callinectes sapidus*, in the Mediterranean Sea, using 22 initial eDNA environmental samples (eDNA) collected from three coastal lagoons.
3. A subsequent large-scale eDNA sampling campaign (61 samples in 31 sites), conducted in collaboration with local stakeholders, was carried out to map the distribution of *Callinectes sapidus* along the Occitanie coastline (Western Mediterranean, France).
4. Using eDNA probe-based qPCR, *Callinectes sapidus* was detected in 32 out the 61 samples (52%), confirming its presence in 24 out of 31 sites surveyed, including the 13 lagoons where its occurrence had already been reported, as well as two additional lagoons and at sea where no prior records existed.
5. Our results demonstrate the utility of eDNA probe-based qPCR for effective monitoring the invasive Blue Crab. The integration of eDNA analysis with citizen science observations enhances the monitoring framework, facilitating early detection and contributing to improved management strategies at the very beginning of species colonization when practical actions could be implemented.

## Introduction

Species that have been introduced to areas outside their natural ranges, known as non-native species, pose a major threat to marine biodiversity with notable socio-economic impacts once they become invasive (Alidoost Salimi et al., 2021; Giakoumi et al., 2019). The introduction of non-native species, and specifically invertebrates, has become a recurring phenomenon over the last century, largely attributed to globalization, international trade (Bailey et al., 2020; Seebens et al., 2021; Zenetos et al., 2022), and has been facilitated by climate change and maritime traffic (Pearman et al., 2021; Seebens et al., 2016). The arrival of these invasive non-native species can significantly jeopardize local biodiversity through phenomena such as competitive exclusion, predation, introgression or niche modification (Mooney & Cleland, 2001). Prevention is widely regarded as the most effective strategy for managing biological invasions, primarily through the establishment of monitoring programs for early detection (Keller et al. 2008). It is generally more cost-effective than controlling or eradicating them post-establishment and reduces the risk of ecological damage, including biodiversity loss and disruption of ecosystem functions. Early detection allows for rapid response measures that can limit the spread and impact of invasive species. Moreover, preventive strategies also provide long-term benefits by reducing future management needs and are often reinforced by public awareness campaigns that enhance community engagement and support for conservation (Giakoumi et al., 2019). Some invasive species have caused significant economic losses by damaging fishing gear and reducing the quality and quantity of commercial catches. In response, artisanal small-scale fishermen have adopted various strategies — as observed in the case of *Portunus segnis* in Tunisia — including changes in net types, reduced setting times, and alternative fishing techniques. However, these adaptations are often costly and labor-intensive (Marchessaux et al., 2023).

The Blue Crab, *Callinectes sapidus* Rathbun, 1896, is indigenous to the estuaries and coastal waters of the western Atlantic coast (USA) where it is actively targeted by fisheries. Its presence in the Mediterranean Sea has been documented as early as 1930 (Galil et al. 2002, Lee et al., 2020). Recent scientific publications confirmed sightings of *C. sapidus* in various locations across the Mediterranean Sea, including Spain, Algeria, Italy and France (Labrune et al. 2019, Mancinelli et al., 2021; Sabelli, 2023). Consequently, the French government has initiated a regional action plan (http://www.occitanie.developpement-durable.gouv.fr/presentation-a25426.html) to provide insights pertinent to the inquiries outlined in European directives concerning lagoon, coastal, and marine environments. As of 2022, Europe has strictly regulated 88 species, including 10 species of fish, 1 species of mollusk and 8 species of aquatic invertebrates including the Chinese mitten crab (*Eriocheir sinensis*, https://circabc.europa.eu/ui/group/4cd6cb36-b0f1-4db4-915e-65cd29067f49/library/79885406-e439-4961-ab2e-717191190f34/details) while *C. sapidus* is in the process of being regulated. One first preventive action is the early detection of this species. Indeed, most reports of the presence of the species in French West Mediterranean Sea are made by fishermen whose nets are often damaged by *C. sapidus* in spring and summer - yet not all lagoons are surveyed, concentration are not the same in the different surveyed lagoons, there are no data for a large part of the potential habitat of *C. sapidus* while the question of its presence in lagoons in winter remains open.

Environmental DNA (eDNA) emerges as a promising tool, enabling early detection of invasive species in low-impacted sites or monitor their persistence in managed sites (Dejean et al., 2012; Kress et al., 2015; Taberlet et al., 2018). Specifically, quantitative polymerase chain reaction (qPCR)-based eDNA detection assays aim to identify a target species through the analysis of DNA present in water samples collected from target sites. Additionally, it can potentially allow for the estimation of a minimum density threshold for species detection (Klymus et al., 2020b, 2020a). The use of qPCR-based approach instead of the eDNA metabarcoding approach has been shown to enhance detection capabilities (McColl-Gausden et al., 2023; Tsuji et al., 2024; Yu et al., 2022). Inventories based on eDNA are particularly interesting in challenging environments such as turbid wetlands colonized by *Callinectes sapidus* (Saenz-Agudelo et al., 2022). Here, we first tested the specificity of a mitochondrial genetic marker to detect the Blue Crab from environmental samples using qPCR-based eDNA detection assays. We then applied qPCR-based eDNA detection assays to monitor specifically the potential presence of C. sapidus across 31 sites comprising 17 lagoons, one channel between lagoons and the sea, three river mouths, one channel between lagoons, and nearshore along the Occitanie coast (French West Mediterranean Sea). Comparing our eDNA-based data with citizen science data and the literature, we discuss how combining such data can improve the monitoring of the invasive Blue Crab.

## Material and Methods

### Study area, eDNA sampling and extraction

Sampling was done along the French Mediterranean coasts following the protocol of Boulanger et al. (2021) (Figure 1).

**Figure 1:**
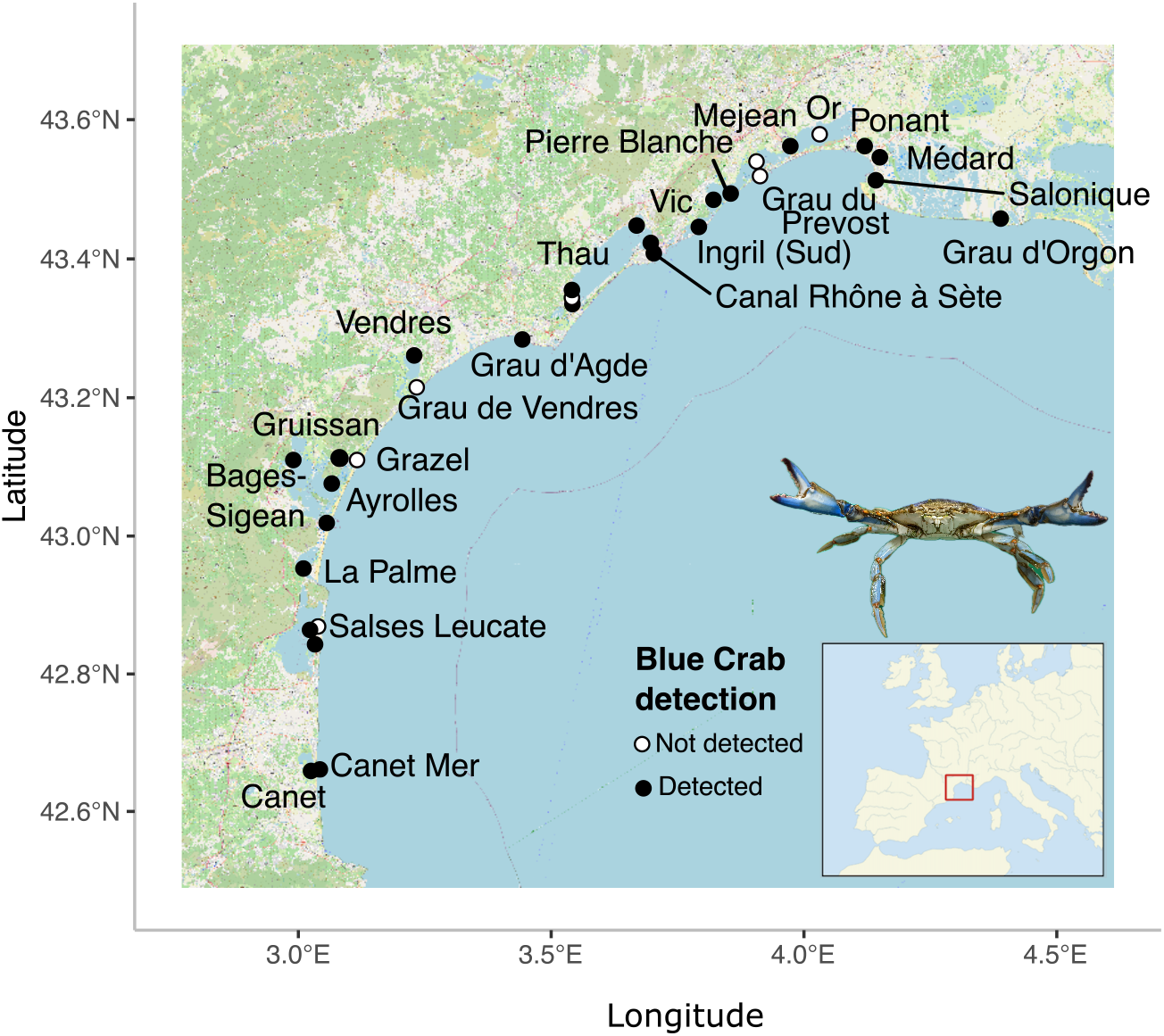
Sampling sites for the eDNA assay targeting the Blue Crab *Callinectes sapidus*. The species is recorded as present when its DNA has been detected in at least one of the two samples collected in each site.

Each water eDNA sample consisted of 30 liters of seawater filtered using a 0.20*µ*M filtration capsule over a 30-minute timed transect from a boat, using a 12 V peristaltic pump with a flow rate of one liter per minute. Immediately after filtration, the capsule was drained of any remaining water, filled with 80 mL of CL1 conservation buffer, and stored at room temperature until extraction. Three sampling campaigns were conducted in collaboration with regional stakeholders with different objectives:

The first campaign to test the ability of our genetic marker to detect *Callinectes sapidus* through water filtration and our protocol of qPCR based eDNA assay, a total of 13 water eDNA samples were collected in situ and in an aquarium (Table 1). Two to three replicates of water samples were performed per site. Sampling in situ were conducted in two sites where a high concentration of *C. sapidus* had been reported by professional fishermen (Canet and Thau lagoons) and at two periods: early in the year in March 2021 at the end of winter when *C. sapidus* is potentially dormant and later during summing (June-July 2021) when the species is active and more abundant. Two additional samples were collected from a 1500L tank hosting around forty *C. sapidus* individuals at the Biodiversarium (Banyuls-sur-Mer) in July 2021 (Table 1).

**Table 1:** Preliminary test from samples collected in 2021 (first campaign).

| Samples | Sites | Date | qPCR |
| --- | --- | --- | --- |
| First campaign |  |  |  |
| SPY210462 | Thau | 23/03/2021 | 7/12 |
| SPY210463 | Thau | 23/03/2021 | 5/12 |
| SPY210459 | Thau | 23/03/2021 | 7/12 |
| SPY210454 | Canet | 24/03/2021 | 4/12 |
| SPY210457 | Canet | 24/03/2021 | 9/12 |
| SPY210453 | Canet | 24/03/2021 | 7/12 |
| SPY210455 | Thau | 01/06/2021 | 9/12 |
| SPY210458 | Thau | 01/06/2021 | 7/12 |
| SPY211260 | Thau | 01/06/2021 | 4/12 |
| SPY212706 | Canet | 27/07/2021 | 5/24 |
| SPY 212707 | Canet | 27/07/2021 | 16/24 |
| SPY 212708 | Biodiversarium | 29/07/2021 | 24/24 |
| SPY 212702 | Biodiversarium | 29/07/2021 | 24/24 |

The second campaign to assess the influence of Blue Crab density on detection, a total of 9 water eDNA samples were collected in July 2022 in three sites with different densities of *Callinectes sapidus*. In each site, three replicates of water samples were collected (Table 2).

**Table 2:** Preliminary test from samples collected in 2022 (second campaign). Sites are categorized by Blue Crab density as follows: high (+++), medium (++), and low (+).

| Samples | Sites | Date | qPCR |
| --- | --- | --- | --- |
| Second campaign |  |  |  |
| SPY221732 | Canet ( +++ ) | 13/07/2022 | 16/17 |
| SPY221731 | Canet ( +++ ) | 13/07/2022 | 0/24 |
| SPY221738 | Canet ( +++ ) | 13/07/2022 | 4/24 |
| SPY221729 | Canet (Roselière) ( ++ ) | 13/07/2022 | 6/24 |
| SPY221737 | Canet (Roselière) ( ++ ) | 13/07/2022 | 0/24 |
| SPY221735 | Canet (Roselière) ( ++ ) | 13/07/2022 | 2/24 |
| SPY221736 | Méjean-Perols ( + ) | 22/07/2022 | 3/24 |
| SPY221734 | Méjean-Perols ( + ) | 22/07/2022 | 1/24 |
| SPY221733 | Méjean-Perols ( + ) | 22/07/2022 | 4/24 |

Sites were chosen in concertation with professional fishermen and stakeholder based, with two sites in Canet and one site in Méjean-Perols. In Canet, the three replicates collected nearby the channel between the lagoon and the sea were considered as the high-density site as the Blue Crab was often caught in fishing nets with high densities. A second site within the same lagoon (Roselière) corresponded to a lower density site of Blue Crab (lower capture rate). Finally, the three replicates collected in Méjean-Perols lagoon also corresponded to a low-density sample site with only episodic captures by professional fishermen.

The third campaign to spatially monitor the presence of *Callinectes sapidus* along the French West Mediterranean Sea coast, we sampled 31 sites from May to June 2023. Two replicates of water samples except for one site (only one water sample) were collected each time, representing a total of 61 eDNA samples (Figure 1, Table S1). Overall, we sampled a diversity of coastal habitats, including (i)17 lagoons: Ayrolles (1 site), Bages-Sigean (2 sites), Canet (1 site), Grazel (1 site), Gruissan (1 site), Ingril (Sud) (1 site), La Palme (1 site), Médard (1 site), Méjean (3 sites), Or (1 site), Pierre Blanche (1 site), Ponant (1 site), Salses-Leucate (3 sites), Salonique (1 site), Thau (4 sites), Vendres (1 site), and Vic (1 site) (ii) one channel between lagoons and the sea: Grau du Prévost (1 site)), (iii) three river mouths: Grau d’Agde (river mouth of the Hérault river, 1 site), Grau de Vendres (river mouth of the Aude river, 1 site) and Grau d’Orgon (river mouth of the Petit Rhône river, 1 site), (iv) one channel between lagoons: Canal du Rhône à Sète (1 site) and (v) nearshore: Canet Mer (1 site). Given the size of some lagoons, we sampled up to four sites within a single lagoon (e.g. Thau). Overall, our study includes not only sites that are surveyed by professional sentinel fishermen where *C. sapidus* had already been reported such as Canet but also sites where no data were available at sampling time: Salonique lagoon, the channel between lagoons, the three river mouths and the channel between lagoon and the sea. *Callinectes sapidus* is suspected to be able to use these accesses to reach the sea or move from one lagoon to another.

### qPCR-based eDNA detection assays

The DNA extraction of eDNA samples was performed at SPYGEN (Le Bourget du Lac, France), following the protocol published by Pont et al. (2018) for the first and the third campaign and in house in a dedicated room for water DNA sample extraction at CEFE for the second campaign (Faure et al., 2023). Some samples of the first campaign extracted at SPYGEN (Le Bourget du Lac, France) have also been re-extracted at CEFE to compare and validate the in-house extraction protocol. Negative extraction controls were carried out in parallel with each batch of eDNA extraction to monitor for potential contamination. Amplifications of the 83 eDNA samples were performed using qPCR following Faure et al., (2023). We used primers and PCR-probe designed by Andersen et al (2018), developed from *Callinectes sapidus* tissue samples in the North Sea to amplify a portion of the mitochondrial cytochrome oxidase 1 genome (see Table S2).

We first verified the specificity of this barcode performing in silico PCR using the online tool Primer-BLAST, with a maximum of 3 mismatches (https://www.ncbi.nlm.nih.gov/tools/primer-blast/index.cgi). We then tested in vivo this barcode on Blue Crab tissue and on six related species found in Canet Lagoon (*Gammarus aequicauda, Palaemon elegans, Eriphia verrucosa, Palaemon adspersus, Idotea chelipes, Palaemon longirostris*), which could potentially confound our qPCR-based eDNA detection assays.

The qPCR-based eDNA detection assays were designed to selectively amplify the DNA of *Callinectes sapidus* and quantify the DNA concentration. An initial calibration step was performed to associate the number of cycles required to reach the exponential phase of DNA amplification, ie cycle threshold (CT) values with a DNA concentration. The qPCR was then applied to each eDNA sample to generate CT values, which were compared to the initial calibration curve to determine the initial DNA quantity. The qPCR runs were performed on the CeMEB labex high-throughput qPCR platform (University of Montpellier), a facility physically separated from the preqPCR laboratory (GEMEX platform, CEFE). We followed Faure et al. (2023) protocol for the qPCR amplification: the final reaction volume of 25 *µ*l contained 3 *µ*l of template DNA (eDNA extract), 12.5 *µ*l of TaqMan™ Environmental Master Mix 2.0 (Life Technologies, now ThermoFisher Scientific), 6.5 *µ*l of double-distilled water (ddH2O), 1 *µ*l of each primer (10 *µ*M) and 1 *µ*l of probe (2.5 *µ*M). Samples were run on a LightCycler® 480 qPCR system (Roche, Basel, Switzerland) under the following thermal cycling conditions: 5 min at 50°C and 10 min at 95°C, followed by 55 cycles of 30 s at 95°C and 1 min at 62°C. The first campaign (2021) was performed in sites with high density of Blue crab and 12 to 24 qPCR replicates were performed for each sample. Following the first results with detections as low as only 5 of 24 replicates positives, we opted for 24 qPCR replicates for all other samples to optimize the early detection of *C. sapidus*. Finally, we sequenced two eDNA qPCR products that were positive under our detection assay to confirm that only Blue Crab had been amplified.

### Citizen science data

The citizen data consisted primarily of reports from professional fishermen, which was then relayed by the Regional Committee of Maritime Fisheries and Marine Farming of Occitanie (CRPMEMO) into the Information System established by Conservatoire d’Espaces Naturels d’Occitanie (SICEN). This information system is a set of resources and devices for collecting, storing, processing and disseminating naturalist data, useful to standardize the data, whether it comes from citizen science or naturalist studies conducted on natural sites. The data is then transferred to the Occitanie SINP (Information System for the Inventory of Natural Heritage). For any new participant in the citizen science program, their observations were first validated by DREAL and CEN Occitanie before being added to the database, provided that the species’ presence was confirmed by photographic evidence. The capture data reported from professional fishermen were considered as already verified observations of *Callinectes sapidus* given their expertise. Overall, observation considered here are the one made between July 2018 and June 2023 for the different lagoons investigated here using qPCR-based eDNA detection assays, providing contemporary information on Blue Crab presence at the time of our sampling in 2023.

## Results

### Sensitivity of the qPCR assay

Two standard curves were produced along the qPCR assay for this study. The first standard curve (y = 3.40x +23,45, R2 = 0.98, efficiency = 97%) has been used for analyzing eDNA samples from the first and second campaign (2023). DNA extracted from Blue Crab tissues was successfully amplified for a range of dilutions with concentrations as low as 6×10-5 ng.*µ*L^-1^, representing the limit of quantification (LOQ). As stock of dilution had been totally used, a second standard curve was produced for the third sampling campaign samples qPCR tests (y = 5.03 x +25.60, R2 = 0.90, efficiency = 58.1%). DNA extracted from Blue Crab tissues was amplified for a range of dilutions with concentrations as low as 4×10-4 ng.*µ*L-1 (LOQ), an order of magnitude higher than for the first standard curve representing the limit of quantification.

### Specificity of the qPCR assay

Positive controls performed on DNA extracted from *Callinectes sapidus* tissues samples displayed the expected DNA concentration values, thereby validating the qPCR assays. None of the six species (tissue) present in the Canet lagoon with Blue Crab were amplified when using the qPCR conditions selected for our test, indicating selectivity of our marker. All eDNA samples from the first campaign, collected in March (late winter) and in June/July (summer) in two lagoons with high densities of Blue Crab, and in aquariums, revealed positive detection of *C. sapidus* (Table 1). We retrieved between 20% (5 positive wells out of 24 for Canet) and 75% of positive wells (9 out of 12 wells for Grau de Canet) in natural habitat and 100% of positive wells for aquariums (24 positive wells out of 24). Sanger sequencing of the qPCR products from two eDNA samples collected in the Biodiversarium (positive control, SPY 212708, SPY 212702) and from two eDNA samples collected in Canet lagoon (SPY212706, SPY212707) (Table 1) matched at 100% with existing Blue Crab sequences already available in NCBI database and confirmed the specificity of our qPCR assay.

### qPCR and effect of Blue Crab density

*Callinectes sapidus* was detected in all three lagoons surveyed during the second field campaign, with seven out of nine samples positive. At both the high- and intermediate-density sites, the species was detected in 2 out of 3 water samples. At the low-density site, all 3 samples were positive.

At the high-density site (Canet), *C. sapidus* DNA was detected in 2 of the 3 filters, with 0%, 16% and 94% of wells testing positive across the three replicates. When detected, CTs values ranged from 36.457 ± 0.57 to 39.05 ± 1.033, corresponding to DNA concentrations between 3.5 E-4 ± 1.5 E-4 to 6.65 E-5 ± 3.34 E-5 ng/µL. At the intermediate-density site, DNA was also detected in 2 out of the 3 filters, with 0%, 8 and 25% of wells positive. CT values ranged from 39.99 ± 0.55 to 40.72 ± 0.165, corresponding to concentrations between 1.86 E-5 ± 2.07 E-6 to 3.18 E-5 ± 1.05 E-5 to ng/µL. Finally, at the low-density site, *C. sapidus* was detected in all three filters, with 4% to 16% of positive wells testing positive. CT values ranged broadly from 25.77 ± 0.28 to 40.35 ± 0.89, corresponding to concentrations from 2.72 E-5 ± 1.78 E-5 to 0.47 ± 0.09.

Overall, the lowest eDNA concentration was observed in the intermediate-density site (1.86 E-5 ± 2.07 E-6) while the highest was recorded at the low-density site. These results suggest that eDNA concentration did not be correlated with estimated crab densities (Table 2). When pooling all wells by site, 31% were positive at the high-density site (Canet) compared to just 11% at both the intermediate- and low-density sites. During the first sampling campaign in 2021, 56% of wells were positive at Canet in March and 44% in July. By contrast, in the most recent campaign (June 2023), only 6% of wells were at that site.

### Comprehensive mapping of Blue Crab distribution in 2023

During each qPCR run, all technical blank controls yielded negative results. A total of 32 eDNA samples (52%) exhibited positive genetic detection in 2023 (Figure 1, Table S1). Overall, *Callinectes sapidus* was detected at 24 sites. At 8 of these sites, both replicates tested positive, while at the remaining site, only one of the two replicates showed detection. No detection was recorded at 7 sites (Or, Méjean Liaison Lez, Grau du Prévost, 1 site at Thau, Grau de Vendres, Grazel, 1 site at Salses-Leucate). We detected *C. sapidus* in all the studied ecosystems, i.e. lagoons, channels between lagoons and sea, river mouths and nearshore. We found that *C. sapidus* was present in 16 out of the 17 lagoons sampled. *Callinectes sapidus* was not detected during this 2023 survey solely in Or lagoon. In large lagoons like Thau and Salses-Leucate, *C. sapidus* has been detected in some but not all sites within each lagoon, highlighting spatial heterogeneity in detection. Notably, while *C. sapidus* was detected in 2022 at both sites in Or lagoon, it was not detected in 2023. *Callinectes sapidus* was consistently detected in Canet, with detections three years in a row (2021, 2022, 2023). No detection occurred in Grau du Prévost, the site representing the channel connecting a lagoon to the sea. The species was detected in two out of the three river mouths surveyed - Grau d’Agde and Grau d’Orgon - but not in Grau de Vendres. Finally, *C. sapidus* has also been detected in Canal du Rhône à Sète which connects several lagoons where *C. sapidus* has been detected in this study, and nearshore in front of Canet lagoon (Canet Mer).

### Comparison of eDNA-based detection with previous records and citizen data

We detected *Callinectes sapidus* in all 10 lagoons where its presence had previously been reported in the literature (Labrune et al 2019, Figure 2). Between July 2018 and June 2023, a total of 181 Blue Crab observations were reported by citizens along the Occitanie coast, 97% of which were originated from professional fishermen. Compared to the literature, citizen science data additionally documented for the first time Blue Crab presence in two of the river mouths we prospected (Grau d’Orgon and Agde) and in three other lagoons (Médard, Vic, Ingril (Sud) ; Figure 2). Our eDNA probe-based qPCR assay also detected *C. sapidus* at each of these five sites. Finally, our eDNA probe-based qPCR assay detected for the first time *C. sapidus* in three sites; two lagoons (Salonique and Pierre Blanche) and nearshore at sea in front of Canet (Figure 2). Notably, no professional fishermen operate in Salonique and Pierre Blanche, which may explain the absence of previous reports. The Blue Crab had already been reported at sea off the coast of Port La Nouvelle.

**Figure 2:**
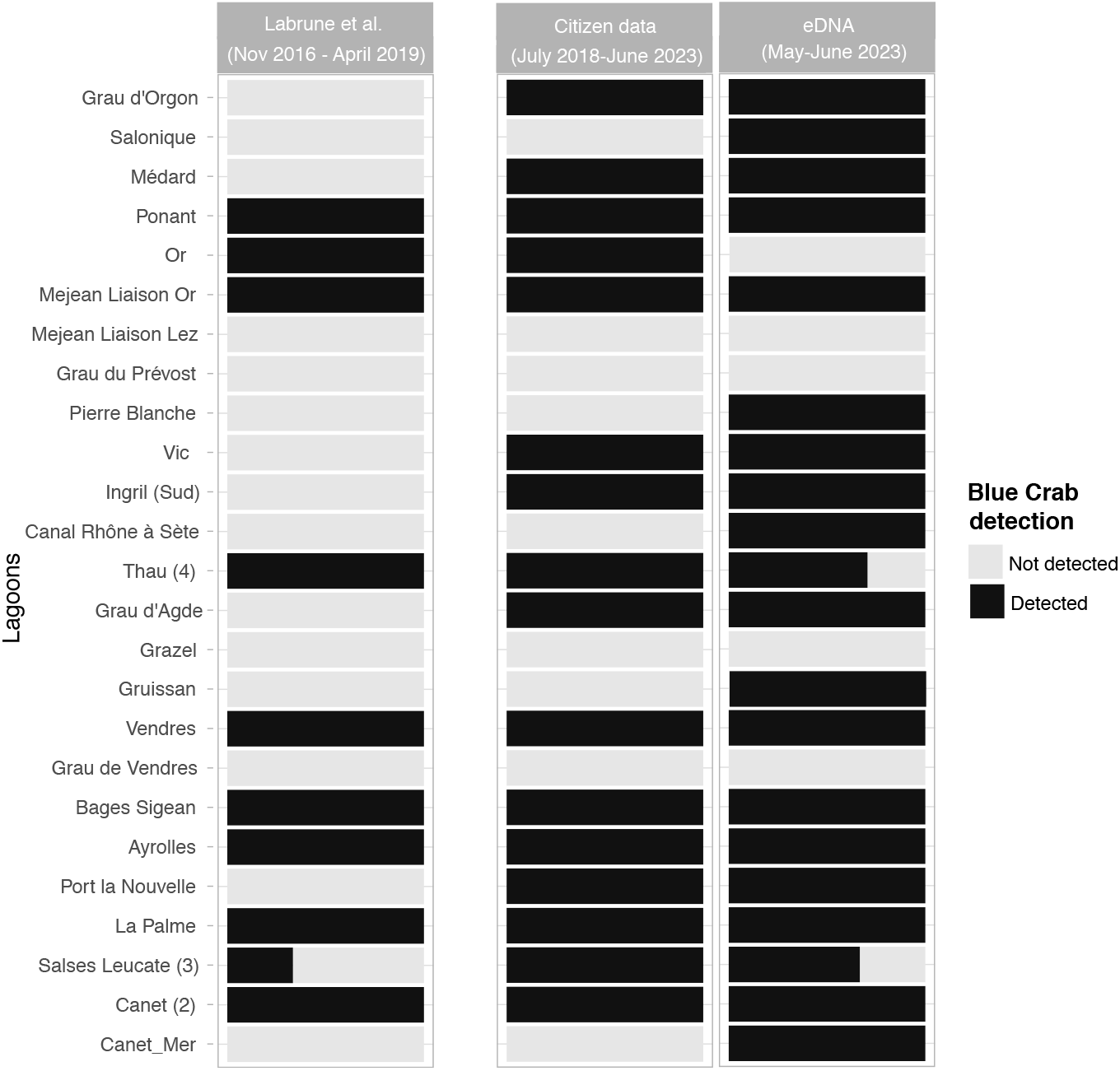
Comparison of the Blue Crab *Callinectes sapidus* detection for all sites sampled and tested with qPCR-based eDNA detection assays (2022; 2 sites and 2023; 31 sites) and comparison with current citizen science data (December 2023) (Stakeholders) and previous records up to 2018 (Labrune et al. 2019). The presence of *Callinectes sapidus* (in black) is reported for eDNA results when its DNA has been detected in at least one of the two samples of a site. Numbers in brackets indicate the number of sites considered for a given location.

## Discussion

Our study demonstrates the effectiveness of eDNA probe-based qPCR for monitoring the invasive Blue Crab *Callinectes sapidus* in lagoons and nearshore in the Mediterranean Sea. The protocol successfully detected the species in 32 samples out 61 tested, confirming its presence in 24 out of the 31 sampled sites. Notably, our eDNA probe-based qPCR tests revealed the presence of *C. sapidus* in the 13 lagoons where the species had been previously documented through independent citizen observations, as well as in three additional sites where no prior records existed at the time of sampling. This study highlights the significant potential of eDNA-based methods for the Blue Crab and more generally invasive species detection and showcases the value of integrating eDNA monitoring with existing databases derived from citizen observations for comprehensive conservation management. Previous studies (Labrune et al. 2019) and citizen observations have already indicated the expansion of *C. sapidus* in the study area (Occitanie), highlighting the necessity for early detection and the urgency of management strategies to mitigate the impact of this invasive species. Our findings not only confirm the presence of *C. sapidus* at sites previously identified by stakeholders and citizen observers but also document new occurrences, demonstrating the high efficacy of eDNA based methods for monitoring invasions in such a context. The utility of eDNA has been established for other invasive invertebrate species in lagoons (Klymus et al., 2017; Roux et al., 2020; Tréguier et al., 2014) and in marine ecosystems (Gargan et al., 2022; Simpson et al., 2023), where it has showed significant advantages over conventional survey methods. For example, the detection rate (per site) of an elusive freshwater turtle species using eDNA qPCR barcoding was 5.55 times higher than conventional trapping methods and the approach proved to be 18.7% more cost effective (Sternhagen et al., 2024). Given this recognized utility, a study developed specific eDNA qPCR assays to detect 60 invasive, threatened, and exploited freshwater vertebrates and invertebrates in Eastern Canada (Hernandez et al., 2020). Given the difficulty of observing the Blue Crab using traditional methods, especially at low density, eDNA qPCR barcoding offers a powerful tool for monitoring the presence of this species.

In our study, eDNA did not enable the quantification of *Callinectes sapidus* populations, as no correlation was found between eDNA concentration and the known density of *C. sapidus*. For fishes, such relationships have been established for some fish species (Karlsson et al., 2022; Rourke et al., 2022) but is not universally (Rourke et al., 2023). For invertebrates, the relation appear less consistent, since they have typically lower shedding rates compared to fish (Allan et al., 2021). Additional factors, such as reproductive behaviors, density or periods of dormancy could further explain the lack of correlation between eDNA concentration and crab density. For instance, Crane et al. (2021) examined the effectiveness of eDNA for detecting the invasive European green crab (*Carcinus maenas*) in North American estuarine environments through both field and laboratory experiments. They found that life stage significantly impacts detection rates, with ovigerous females showing the highest eDNA concentrations. Similarly, Marques et al. (2024) developed and tested an eDNA assay to detect the critically endangered fan mussel (*Pinna nobilis*) and found that although overall eDNA concentrations were low, sampling depth near the seafloor and the occurrence of a putative reproductive event significantly increased eDNA detection. An eDNA assay has also been used to monitor populations of the Crown-of-Thorns Seastar (*Acanthaster* cf. *solaris*), a significant coral predator, on the Great Barrier Reef, to assess how eDNA detection correlates with actual Crown-of-Thorns Seastar densities (Uthicke et al., 2022). The authors reported that while eDNA copy numbers align with higher Crown-of-Thorns Seastar densities, the relationship became much less reliable at lower, sub outbreak densities. In our study, we observed substantial variability between replicates of water samples, with most detection occurring in only one of the two replicates. This variability may help explain the lack of correlation and underscores the importance of using multiple replicates per site. Finally, our first sampling campaign demonstrated that *C. sapidus* can be detected even during periods when the species is presumed to be dormant. We sampled at the end of winter, when activity is expected to be low, and again in summer, when the species is typically more active and abundant in the two lagoons. Remarkably, *C. sapidus* was detected under both conditions. However, a proper assessment of eDNA detectability during dormancy will require a targeted study using a dedicated sampling and validation protocol.

Combining both eDNA and citizen/stakeholder observations has proven to be an effective way to enhance avian ecological research (Padró, 2024). Here, we show that it offers a dual advantage. First it provides an opportunity to validate the eDNA as an adequate tool for detecting *Callinectes sapidus* populations. Second, the two approaches are complementary. Citizen and stakeholders-based monitoring may be readily accessible in some sites while it may prove to be more challenging in others where eDNA provides an effective alternative. While observations provide a direct result, eDNA probe-based qPCR approach is faster, cheaper and more sensitive than eDNA metabarcoding approach. Yet, they are still not as fast as direct observations. One key benefit of eDNA is that it enables more systematic and less opportunistic monitoring compared to citizen observations. This facilitates standardized, large-scale spatial monitoring (e.g. Mathon et al., 2023) and/or seasonal or temporal assessments (Bálint et al., 2018). Little is known about their distribution during winter. Citizen and professional fishermen observations primarily detect *C. sapidus* from May to October, when involved people are the more present in the field and seasonal fishing activities occur. Our preliminary results show that the Blue Crab could be detected even during the winter season. Additionally, both citizen and professional fishermen observations have difficulties in detecting larval or juvenile stages (2-5 cm). The development of eRNA is promising, with the potential to identify early life stages and even distinguish between juvenile and adult individuals (Parsley & Goldberg, 2024) is particularly promising. eDNA/RNA can thus be a powerful tool to address these knowledge gaps in future studies. Regarding *C. sapidus* monitoring, channels connecting the sea and lagoons, serve as strategic passageways throughout the year. We recommend prioritizing monitoring in such critical sites to determine their role in the invasion process, either as vectors of dispersal or as refugia during the winter months.

Integrating environmental DNA analysis with targeted barcode-based detection offers a straightforward and reproducible approach for monitoring population fluctuations over time. Here, we presented a protocol specifically designed for detecting the Blue Crab along the French Mediterranean coast. This methodology demonstrated significant potential to support stakeholder engagement in population monitoring by offering an accessible and reliable tool for long-term ecological surveillance and conservation efforts.

## Data Availability Statement

The data that support the findings of this study are available in Supplementary material. Detection of the *Callinectes sapidus* have also been uploaded on the Natural Héritage Inventory Information System (SINP) and on CEN Occitanie Lizmap.

## Ethics and Permit Approval Statement

No ethical nor permit were required for the research should be stated here.

## Acknowledgements

The authors would like thank Stéphane Hourdez (LECOB, Banyuls-sur-Mer) for sharing tissues of invertebrates. The authors would like to address a special thank to all sites’ managers for their help with this study: Canet (Rolland Mivière & Thierry Auga-Bascou; Syndicat mixte du bassin versant du Réart (SMBVR), Office Français de la Biodiversité – Parc Natural Marin du Golfe du Lion (PN-MGL)); Salses-Leucate (Jean-Francois Laffon ; Syndicat mixte RIVAGE, Conseil Départemental des Pyrénées-Orientales) ; La Palme and Bages-Sigean (Parc naturel régional de la Narbonnaise, OFB SD 11) ; Vendres (Rémi Belleza ; Communauté de communes La Domitienne et sa Capitainerie) ; Thau (Michela Patrissi ; Centre d’étude pour la Promotion des Activités Lagunaires et Maritimes (Cépralmar), Syndicat Mixte du Bassin de Thau (SMBT)); Ingril Sud (Romain Nathan Danielou ; Sète Agglopole Méditerranée, Conservatoire du Littoral), Vic and Pierre Blanche (Romain Nathan Danielou CEN Occitanie); Méjean-Pérols et Or (Max Lecouturier ; Ville de Lattes, Syndicat Mixte du Bassin de l’étang de l’Or (SYMBO)); Grau d’Agde (Sylvain Blouet, Florent Keller, Patrick Ramy; Aire Marine de la côte Agathoise); Grau du Prévost (Mondy Laigle; Association de gestion de la Réserve Marine de la côte palavasienne); Ponant, Médard et Salonique (Christophe Rosso ; Mairie du Grau du Roi, Syndicat Mixte pour la protection et la gestion de la Camargue Gardoise (SMCG)); Grau d’Orgon (Delphine Marobin; Parc naturel régional de Camargue). We also thank the numerous students from CEFE and MARBEC labs for their valuable help in the field, including - but not limited to - Dimitri Médétian, Maurine Vilcot, Margot Dentan, Andréa Reyes Camargo, Simon Bettinger, Marie Orblin, and Loïc Sanchez.

## Author contributions

EDT, CM, NB, LC and SM contributed to the study conception and design. All authors were involved in the Material preparation, data collection and analysis. The first draft of the manuscript was written by EDT and SM, and all authors commented on previous versions of the manuscript. All authors have read and approved the final manuscript.

