## Supplementary material for "Improving the monitoring of the invasive Blue Crab (*Callinectes sapidus*): combining environmental DNA and citizen observations": Table S1 and Table S2

**Table S1**: Samples collected in 2023 in collaboration with stakeholders (third campaign).

**Table S2**: Primers and probes specific for *Callinectes sapidus* developed by Andersen et al. (2018), targeting a 275 base pair long fragment from the mitochondrial cytochrome oxidase 1 gene.

**Table S1**

| Date | Site | Type of site | Sample | no. Positive qPCR | mean CT | Mean concentration (ng/µl) | GPS coordinates |
| --- | --- | --- | --- | --- | --- | --- | --- |
| 20230524 | Or #1 | Lagoon | SPY230139 | 0/24 |  |  | 43.58214, 4.01690 |
| 20230524 | Or #2 | Lagoon | SPY230151 | 0/24 |  |  | 43.58214, 4.01690 |
| 20230523 | Ponant #1 | Lagoon | SPY230119 | 15/24 | 39 | 1.19E-05 | 43.56424, 4.11938 |
| 20230523 | Ponant #2 | Lagoon | SPY230120 | 7/24 | 40 | 2,56E-06 | 43.56424, 4.11938 |
| 20230621 | Méjean liaison Or #1 | Lagoon | SPY230108 | 0/24 |  |  | 43.55134, 3.96770 |
| 20230621 | Méjean liaison Or #2 | Lagoon | SPY230116 | 2/24 | 40 | 5.34E-5 | 43.55134, 3.96770 |
| 20230524 | Méjean #1 | Lagoon | SPY230137 | 2/24 | 38 | 2.16E-05 | 43.54771, 3.96134 |
| 20230523 | Médart #1 | Lagoon | SPY230128 | 4/24 | 40 | 4.26E-06 | 43.54379, 4.14117 |
| 20230523 | Médart #2 | Lagoon | SPY230176 | 2/24 | 40 | 8.85E-03 | 43.54379, 4.14117 |
| 20230524 | Méjean liaison Lez #1 | Lagoon | SPY230144 | 0/24 |  |  | 43.53508, 3.91513 |
| 20230524 | Méjean liaison Lez #2 | Lagoon | SPY230152 | 0/24 |  |  | 43.53508, 3.91513 |
| 20230517 | Canal Rhône à Sète #1 | Channel between lagoons | SPY230132 | 2/24 | 40 | 1.14E-06 | 43.53414, 3.91761 |
| 20230517 | Canal Rhône à Sète #2 | Channel between lagoons | SPY230160 | 0/24 |  |  | 43.53414, 3.91761 |
| 20230517 | Grau du Prévost #1 | Channel between lagoon and sea | SPY230154 | 0/24 |  |  | 43.51845, 3.91507 |
| 20230517 | Grau du Prévost #2 | Channel between lagoon and sea | SPY230164 | 0/24 |  |  | 43.51845, 3.91507 |
| 20230523 | Salonique #1 | Lagoon | SPY230127 | 2/24 | 38 | 1.84E-05 | 43.51032, 4.13986 |
| 20230523 | Salonique #2 | Lagoon | SPY230135 | 2/24 | 40 | 1.83E-06 | 43.51032, 4.13986 |
| 20230607 | Vic #1 | Lagoon | SPY230107 | 16/24 | 39 | 1.50E-05 | 43.50387, 3.82136 |
| 20230607 | Vic #2 | Lagoon | SPY230175 | 1/24 | 39 | 1.26E-02 | 43.50387, 3.82136 |
| 20230607 | Pierre Blanche #1 | Lagoon | SPY230161 | 0/24 |  |  | 43.49182, 3.84864 |
| 20230607 | Pierre Blanche #2 | Lagoon | SPY230106 | 15/24 | 40 | 1.20E-05 | 43.49182, 3.84864 |
| 20230524 | Grau d’Orgon #1 | River mouth | SPY230136 | 3/24 | 40 | 4.30E-06 | 43.44517, 4.40013 |
| 20230524 | Grau d’Orgon #2 | River mouth | SPY230172 | 3/24 | 39 | 1.33E-02 | 43.44517, 4.40013 |
| 20230607 | Ingril (Sud) #1 | Lagoon | SPY230105 | 15/24 | 39 | 1.03E-05 | 43.43586, 3.77617 |
| 20230607 | Ingril (Sud) #2 | Lagoon | SPY230178 | 1/24 | 38 | 1.84E-02 | 43.43586, 3.77617 |
| 20230525 | Thau #1 | Lagoon | SPY230121 | 3/24 | 40 | 7.84E-07 | 43.43038, 3.61973 |
| 20230525 | Thau #2 | Lagoon | SPY230165 | 0/24 |  |  | 43.43038, 3.61973 |
| 20230525 | Thau #3 | Lagoon | SPY230129 | 2/24 | 40 | 1.52E-06 | 43.41435, 3.68625 |
| 20230525 | Thau #4 | Lagoon | SPY230166 | 0/24 |  |  | 43.41435, 3.68625 |
| 20230525 | Thau #5 | Lagoon | SPY230138 | 1/24 | 40 | 1.58E-06 | 43.34731, 3.53668 |
| 20230525 | Thau #6 | Lagoon | SPY230145 | 0/24 |  |  | 43.34731, 3.53668 |
| 20230525 | Thau #7 | Lagoon | SPY230153 | 0/24 |  |  | 43.34588, 3.54080 |
| 20230525 | Thau #8 | Lagoon | SPY230168 | 0/24 |  |  | 43.34588, 3.54080 |
| 20230515 | Grau d'Agde #1 | River mouth | SPY230122 | 5/24 | 39 | 2.35E-05 | 43.28384, 3.44399 |
| 20230515 | Grau d'Agde #2 | River mouth | SPY230131 | 0/24 |  |  | 43.28384, 3.44399 |
| 20230509 | Vendres #1 | Lagoon | SPY230124 | 2/24 | 41 | 4.38E-07 | 43.26368, 3.22031 |
| 20230509 | Vendres #2 | Lagoon | SPY230155 | 0/24 |  |  | 43.26368, 3.22031 |
| 20230509 | Grau de Vendres #1 | River mouth | SPY230150 | 0/24 |  |  | 43.21638, 3.23452 |
| 20230509 | Grau de Vendres #2 | River mouth | SPY230158 | 0/24 |  |  | 43.21638, 3.23452 |
| 20230509 | Gruissan #1 | Lagoon | SPY230134 | 1/24 | 41 | 2.19E-07 | 43.11652, 3.08581 |
| 20230509 | Gruissan #2 | Lagoon | SPY230141 | 0/24 |  |  | 43.11652, 3.08581 |
| 20230509 | Grazel #1 | Lagoon | SPY230142 | 0/24 |  |  | 43.11352, 3.11834 |
| 20230509 | Grazel #2 | Lagoon | SPY230156 | 0/24 |  |  | 43.11352, 3.11834 |
| 20230531 | Bages-Sigean #1 | Lagoon | SPY230114 | 15/24 | 40 | 1.10E-05 | 43.11247, 2.99626 |
| 20230531 | Bages-Sigean #2 | Lagoon | SPY230159 | 0/24 |  |  | 43.11247, 2.99626 |
| 20230510 | Ayrolles #1 | Lagoon | SPY230123 | 4/24 | 42 | 1.02E-06 | 43.07572, 3.07572 |
| 20230510 | Ayrolles #2 | Lagoon | SPY230147 | 0/24 |  |  | 43.07572, 3.07572 |
| 20230531 | Bages-Sigean #3 | Lagoon | SPY230146 | 0/24 |  |  | 43.03522, 3.02935 |
| 20230531 | Bages-Sigean #4 | Lagoon | SPY230115 | 20/24 | 39 | 2.34E-05 | 43.03522, 3.02935 |
| 20230510 | La Palme #1 | Lagoon | SPY230133 | 1/24 | 41 | 2.67E-07 | 42.96749, 3.00029 |
| 20230510 | La Palme #2 | Lagoon | SPY230148 | 0/24 |  |  | 42.96749, 3.00029 |
| 20230510 | Salses-Leucate #1 | Lagoon | SPY230125 | 5/24 | 40 | 2.80E-06 | 42.86663, 3.02109 |
| 20230510 | Salses-Leucate #2 | Lagoon | SPY230130 | 3/24 | 41 | 6.71E-07 | 42.86663, 3.02109 |
| 20230510 | Salses-Leucate #3 | Lagoon | SPY230140 | 0/24 |  |  | 42.85438, 3.03137 |
| 20230510 | Salses-Leucate #4 | Lagoon | SPY230157 | 0/24 |  |  | 42.85438, 3.03137 |
| 20230510 | Salses-Leucate #5 | Lagoon | SPY230126 | 3/24 | 40 | 2.54E-06 | 42.85295, 3.03234 |
| 20230510 | Salses-Leucate #6 | Lagoon | SPY230149 | 0/24 |  |  | 42.85295, 3.03234 |
| 20230619 | Canet Mer #2 | Coastal site | SPY230174 | 0/24 |  |  | 42.66224, 3.03546 |
| 20230619 | Canet Mer #1 | Coastal site | SPY230177 | 1/24 | 41 | 3.5E-5 | 42.66224, 3.03546 |
| 20230619 | Canet #2 | Lagoon | SPY230109 | 2/24 | 42 | 1.6E-5 | 42.65927, 3.02769 |
| 20230619 | Canet #1 | Lagoon | SPY230113 | 1/24 | 41 | 3.21E-5 | 42.65927, 3.02769 |

**Table S2**

| **Oligo name** | **oligo sequence in 5'->3' direction** | **Temp (°C)** | **Length (bp)** | **GC (%)** |
| --- | --- | --- | --- | --- |
| Calsap_co1_F01 | 5'-GGGCCTCAGTTGATCTTGGT-3' | 59.7 | 20 | 55.0 |
| Calsap_co1_P01 | 5'-FAM-ATACCTCATTCTTCGACCCAGCTGGAG-BHQ1-3' | 59.5 | 20 | 60.0 |
| Calsap_co1_R01 | 5'-GTAGAGAACAGGGTCGCCTC-3' | 65.8 | 27 | 51.9 |
